# Chromatin context-dependent effects of epigenetic drugs on CRISPR-Cas9 editing

**DOI:** 10.1101/2023.05.02.539047

**Authors:** Ruben Schep, Max Trauernicht, Ben Morris, Xabier Vergara, Roderick L. Beijersbergen, Bas van Steensel

**Affiliations:** Oncode Institute, The Netherlands; Division of Gene Regulation, Netherlands Cancer Institute, 1066 CX Amsterdam, The Netherlands; Division of Molecular Carcinogenesis, Netherlands Cancer Institute, 1066 CX Amsterdam, The Netherlands; Division of Cell Biology, Netherlands Cancer Institute, 1066 CX Amsterdam, The Netherlands

**Keywords:** CRISPR, chromatin, drug screen, HDAC, Double-stranded break repair, Decitabine

## Abstract

The efficiency and outcome of CRISPR/Cas9 editing depends on the chromatin state at the cut site. It has been shown that changing the chromatin state can influence both the efficiency and repair outcome, and epigenetic drugs have been used to improve Cas9 editing. However, because the target proteins of these drugs are not homogeneously distributed across the genome, the efficacy of these drugs may be expected to vary from locus to locus. Here, we systematically analyzed this chromatin context-dependency for 160 epigenetic drugs. We used a human cell line with 19 stably integrated reporters to induce a double-stranded break (DSB) in different chromatin environments. We then measure Cas9 editing efficiency and repair pathway usage by sequencing the mutational signatures. We identified 67 drugs that modulate Cas9 editing efficiency and/or repair outcome dependent on the local chromatin environment. For example, we find a subset of histone deacetylase inhibitors that improve Cas9 editing efficiency throughout all types of heterochromatin (e.g., PCI-24781), while others were only effective in H3K27me3-marked regions (e.g., Vorinostat). In summary, this study reveals that most epigenetic drugs alter CRISPR editing in a chromatin-dependent manner, and provides a detailed guide to improve Cas9 editing more selectively at the desired location.

**HIGHLIGHTS:**

- A screen identifies dozens of drugs that alter Cas9 editing in a chromatin context-dependent manner
- Many HDAC inhibitors boost Cas9 editing efficiency throughout all types of heterochromatin
- The DNMT inhibitor Decitabine completely blocks resection-dependent repair across the genome

## INTRODUCTION

Cas9-mediated double-strand break (DSB) induction has become an invaluable tool for genome editing (1,2). Cas9 editing in eukaryotic cells has been optimized in various ways (3,4). However, one remaining challenge is that the packaging of DNA into chromatin can hinder Cas9 from binding and cutting DNA (5–10). This is particularly the case in heterochromatin, the compacted form of chromatin (5,11–15). Silenced and cytosine-methylated DNA was also found to be relatively refractory to Cas9 editing (16).

Cas9-induced breaks can be repaired by multiple pathways, including nonhomologous end joining (NHEJ), homologous recombination (HR), and microhomology-mediated end joining (MMEJ, also referred to as theta-mediated end joining) (17,18). Each of these pathways can yield different repair products: MMEJ typically results in small deletions, NHEJ can yield either small insertions or scar-less repair, and HR is thought to be essentially error-free (19–21). Thus, the outcome of a Cas9 editing experiment is strongly dictated by the relative activities of these pathways.

Evidence is accumulating that the local chromatin environment affects this pathway balance. For example, DSBs in heterochromatic regions are more frequently repaired by MMEJ than euchromatic regions (15), while contacts with the nuclear envelope can also alter pathway balance (22). Such effects may at least in part be explained by interactions of pathway-specific repair proteins with certain chromatin features such as histone modifications (23,24). For example, CtIP, an important factor for DNA end resection (a key step in MMEJ and HR), can be specifically recruited to H3K36me3 (25,26). Another example is 53BP1, a key factor in the first steps of NHEJ. This protein is preferentially recruited at sites with H4K20me2 (27–29), but its recruitment is blocked by H4K16 acetylation (30–32).

Together, these findings highlight the importance of the local chromatin context for Cas9 editing, as it affects (1) the editing efficiency and (2) the balance between repair pathways, and hence the probability of obtaining a desired mutation. For this reason, several studies have explored whether chromatin-modifying drugs may be employed to improve Cas9 editing (18,33–36). These studies primarily focused on histone deacetylase (HDAC) inhibitors because these drugs are thought to cause decompaction of chromatin. Some HDAC inhibitors appeared to improve Cas9 genome editing, but to variable degrees (33–36). Importantly, because the binding and activity of HDACs varies across the genome (37), this effect may be expected to depend on the local chromatin context. Indeed, the HDAC inhibitor Trichostatin A was found to improve Cas9 editing in four loci with low levels of H3K27ac, but much less in two loci with high levels of H3K27ac (35). This strongly suggests that chromatin-modifying drugs may act differently across the genome depending on the local chromatin context. However, in the context of Cas9 editing the available evidence is only anecdotal; systematic analysis of this dependency on chromatin context has been lacking.

Besides HDAC inhibitors, a broad diversity of other chromatin-modifying drugs has been developed. Such drugs are candidates to improve Cas9 editing, but no systematic survey has been reported to our knowledge. It is likely that the effects of these drugs are dependent on the local chromatin context, because all these drugs target specific chromatin proteins that are not uniformly active across the genome.

Here, we report a systematic analysis of the chromatin context-dependent effects of 160 chromatin-modifying drugs on Cas9 editing in human cells. For this we used a previously described reporter that tracks the overall editing frequency and relative activities of MMEJ and NHEJ (15). We inserted this reporter into 19 different genomic locations that together represent all major types of chromatin, and then determined the editing efficiency and MMEJ:NHEJ balance in each of these locations after treatment with each of the 160 drugs. We applied a statistical framework to correlate the drug activities with the presence or absence of a wide range of chromatin features. We thus identified many drugs that alter Cas9 editing and show that in many instances the local chromatin context can affect the efficacy of a drug. These results underscore the effect of chromatin on the efficacy of many drugs. Suitable chromatin-modifying drugs can thus be selected to improve genome editing in a genomic locus of interest.

## RESULTS

### Overview of the reporter system and the design of the screen

To assess chromatin context-dependent effects of a large set of drugs on Cas9 editing, we developed a screen based on our sequencing-based reporter assay (15). This assay employs a reporter that carries a short sequence that, when it is cut by Cas9, can be repaired by either NHEJ or MMEJ. These repair pathways predominantly produce signature insertions of one nucleotide (+1) or deletions of seven nucleotides (−7), respectively (20). These signature mutations can be used to infer the relative activity of the two pathways (15,20).

We employed a human K562 cell line (clone #5) that carries 19 randomly integrated copies of this reporter, each marked by a unique 16-nucleotide barcode (15). We refer to these reporters as Integrated Pathway Reporters (IPRs). Clone #5 also carries a cassette for inducible expression of Cas9 (20). Upon activation of Cas9, accumulation of pathway-specific insertions and deletions (indels) can be determined by high-throughput sequencing. The key advantage of this approach is that the repair outcomes can be monitored in parallel in 19 distinct genomic regions, owing to the unique barcode that marks each IPR.

Importantly, the 19 IPRs are located in highly diverse chromatin environments, including transcribed genes and enhancers (euchromatin); H3K27me3-marked heterochromatin; and regions of heterochromatin that interact with the nuclear lamina, replicate late in S-phase, and are marked by H3K9 di- or trimethylation (here referred to as triple heterochromatin) (**Figure 1A**). There are also four IPRs in a mixed state that cannot be easily attributed to any of the other groups. As reported (15) the 19 IPRs display reproducible differences in total indel frequency (TIF) and in the relative abundance of MMEJ and NHEJ (MMEJ:NHEJ ratio, calculated as the ratio between the signature indels), which can be attributed to the different chromatin contexts of each IPRs (**Figure 1B**). Thus, we expected that we should be able to quantitatively detect any effect that a chromatin-modifying drug might have in each of the 19 different contexts (i.e., local effects). Additionally, we can quantify global effects as the average measures of all the IPRs.

**Figure 1.**
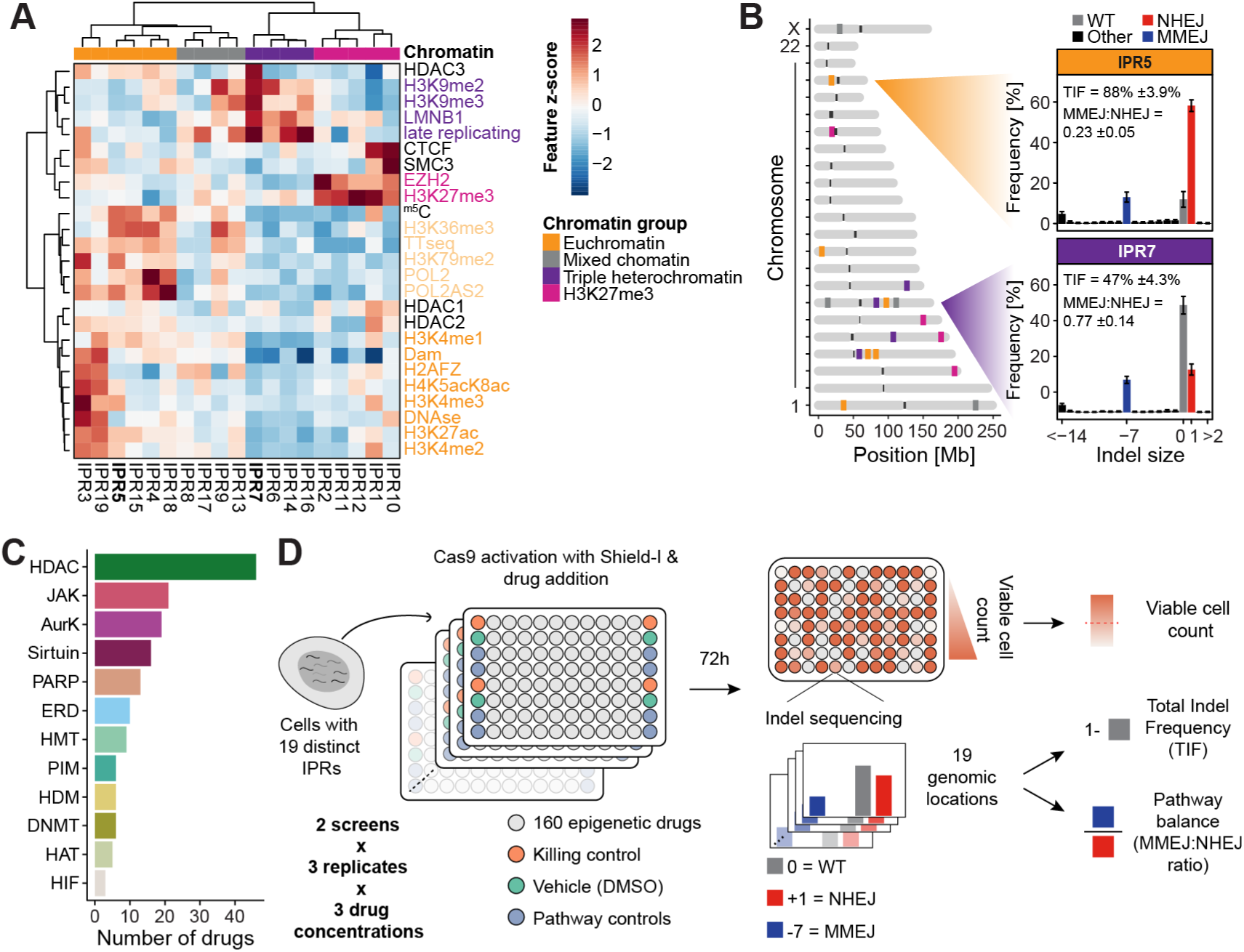
Overview of the reporter system and screen data. (A) Heatmap of the chromatin landscape of the K562 clone #5 used in this study. Z-scores of the 25 chromatin features are indicated of all 19 IPRs. Chromatin features are colored by their type, and IPRs were clustered into chromatin groups based on the chromatin features. IPRs in bold are the two example IPRs in B. (B) Chromosome ideogram with the mapped integrated reporters with an indel plot representing the indels in the control setting for two representative IPRs, one in euchromatin (IPR5), and one in heterochromatin (IPR7). (C) Number of drugs per target group in the screen. HDAC = histone deacetylase, JAK = Janus kinase, AurK = aurora kinase, PARP = poly (ADP-ribose) polymerase, ERD = epigenetic reader domain, HMT = histone methyltransferase, PIM = proviral integration site for moloney murine leukemia virus kinase, HDM = histone demethylase, DNMT = DNA methyltransferase, HAT = histone acetyltransferase and HIF = hypoxia inducible factor (D) Schematic overview of the experimental layout and the readouts of the drug screen.

We assembled a drug library from two commercially available epigenetic drug collections, consisting of 160 drugs in total (**Figure 1C, Table S1**). Among these we included 46 different HDAC inhibitors. Some inhibitors of this class can affect Cas9 editing (33–36), yet HDAC inhibitors greatly vary in their target specificity (e.g.,38). Other tested drugs included inhibitors of Janus kinases (JAK), aurora kinases (AurK), sirtuins, poly (ADP-ribose) polymerase (PARP), epigenetic reader domain-containing proteins (ERD), PIM kinase, histone methyltransferases (HMT), histone acetyltransferases (HAT), DNA methyltransferases (DNMT), histone demethylases (HDM) and hypoxia-inducible factors (HIF). We used a 96-wells automated setup to administer the drug library to clone #5 cells in an arrayed format (**Figure 1D**). Additionally, 24 separate wells received the vehicle (DMSO solvent) only, serving as negative controls. We also included treatment with Mirin (an inhibitor of MMEJ) and NU7441 (an inhibitor of NHEJ) as positive controls. Simultaneously, we activated Cas9 by inhibition of its degron-mediated degradation (39) and transfected a single-guide RNA (sgRNA) that targets the IPR sequence. After 72 hours we collected genomic DNA and sequenced the IPRs, including their barcode, to quantify the TIF and the MMEJ:NHEJ ratio for each of the 19 IPRs (**Figure 1D**). We also estimated the number of viable cells in each well to monitor any cytotoxic effects that the drugs might have. We performed this screen twice (with an interval of more than a year), each time in three replicates in which the cells were cultured and transfected separately.

Because the 160 drugs used in this screen have a wide range of effective concentrations and may have concentration-dependent toxic side effects, we tested each drug at three different concentrations (100 nM, 1 µM, and 10 µM). We then excluded wells from the analysis that had fewer than 25% viable cells compared to the DMSO treated control in at least two of the three replicates of either screen. Applying this filter, we removed data for 5, 19 and 48 drugs at 100 nM, 1 µM, and 10 µM, respectively (**Figure S1A-D**), while also verifying reproducibility between screens (**Figure S1A**).

Both the TIF and the MMEJ:NHEJ ratio of the vehicle control samples were highly reproducible between the two screens across the 19 IPRs (**Figure S1E-F**). The variation observed in TIF and MMEJ:NHEJ ratio effect size between replicates was likely due to differences in sgRNA transfection efficiency. As previously observed (15), IPRs in euchromatin typically displayed a higher TIF and a lower MMEJ:NHEJ ratio compared to heterochromatic IPRs (**Figure S1E-F**). For downstream statistical analysis we then converted the drug-induced changes in TIF and MMEJ:NHEJ ratio for each IPR to z-scores, using the 24 vehicle control samples per replicate to estimate null distributions (**Figure S1G-H**). The drug-induced changes in TIF were highly reproducible across all replicates and between screens (**Figure S1I**). Changes in MMEJ:NHEJ ratio varied more, potentially due to intrinsic higher variation, as observed in the control condition (**Figure S1J, S1F**). However, after combining the three replicates of each screen, the local (**Figure S1K-L**) and global (**Figure S1M-N**) significant z-scores for both the TIF and MMEJ:NHEJ ratio correlated strongly between the two screens. We conclude that the results of the screens are sufficiently reproducible, and we therefore combined the z-scores of all replicates into one final z-score for each drug-IPR combination, separately for the TIF and MMEJ:NHEJ scores.

### Chromatin-perturbing drugs affecting editing efficiency

We first set out to identify the drugs that induced changes in TIF globally, i.e., on average across all 19 IPRs. In total, 15 (14 increased | 1 decreased), 11 (6|5), and 15 (10|5) drugs significantly changed the TIF globally at 100 nM, 1 µM, and 10 µM, respectively (*p_adj._* < 0.01) (**Figure 2A**). Among the 34 drugs that have a significant global effect at one or more concentrations, 14 are HDAC inhibitors, which is a significant enrichment (Fisher’s exact test *p_adj._* = 9.99 × 10^-3^) (**Figure 2B**). These 14 HDAC inhibitors all caused a global increase in TIF, although several showed a quantitative dependency on chromatin context (discussed below). Of previously reported HDAC inhibitors (33,35), our screen detected PCI-24781 as one of the strongest hits and Trichostatin A (TSA) showed a weaker effect. Our findings indicate that many different HDAC inhibitors, most of which have not been reported before, are capable of increasing genome editing efficiencies.

**Figure 2.**
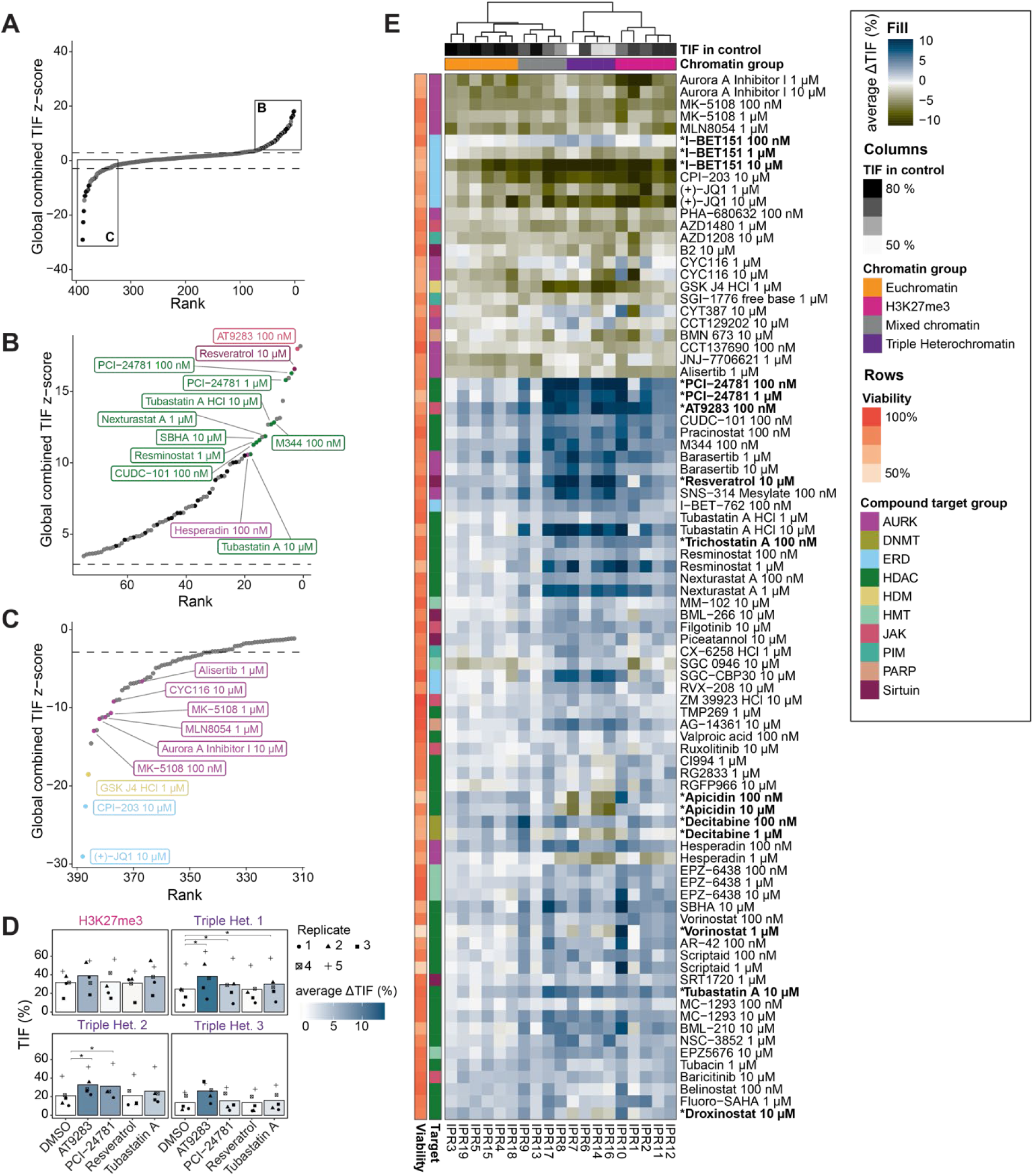
Drugs with strongest effects on TIF. (A) Global (average across all IPRs) change in TIF, ranked from highest increase to strongest decrease. Dashed lines indicate significance threshold of *padj.* < 0.01. Top and bottom 75 hits are zoomed in in B and C respectively. Black dots are samples with a significant global effect in both screens, grey dots are non-significant or only significant in one screen. (B) Overview of the 75 strongest TIF-boosting drugs passing the viable cell count filter. The top 10 significant and reproducible hits are annotated and colored based on their target. Color code in E. (C) Strongest TIF-reducing drugs, included are only drugs with > 70% relative viable cell count. (D) Validation of the strongest TIF-boosting drugs in additional heterochromatic loci. Displayed is the TIF of the four different heterochromatic genomic loci (one H3K27me3-demarcated locus and three triple heterochromatin loci) treated with either vehicle, AT9283 (100 nM), PCI-24781 (100 nM), Resveratrol (10 µM), or Tubastatin A (10 µM). The color gradient indicates the average ΔTIF in percent. The shapes represent the different replicates, the bar indicates the average TIF. TIF were compared between DMSO-treated and drug-treated samples using paired t-tests, with FDR-correction, * = *padj* < 0.05. (E) Heatmap of the local ΔTIF per IPR of all drugs significantly affecting at least one IPR (*padj.* < 0.01). The rows are annotated with the relative viable cell count and drug target. The column annotation contains the chromatin domain and TIF in the control. The rows were clustered based on the average effect of the concentrations of the drug. Drugs discussed in the text are highlighted in bold.

Besides HDAC inhibitors, several other drugs, for which the potential to increase editing efficiency has not been reported before, globally enhanced TIF. Among the strongest hits are Resveratrol (a Sirtuin inhibitor), AT9283 (an inhibitor of both JAK2/3 and Aurora kinases) and Hesperadin (another Aurora kinase inhibitor). These data suggest that a diversity of chromatin-modifying drugs may be used to improve CRISPR editing.

We further examined the effects of four of the strongest TIF-increasing drugs (AT9283, Resveratrol, PCI-24781, Tubastatin A) in four native genomic sequences (*i.e.*, outside the 19 IPRs) in heterochromatic loci (**Figure S2C**). We used TIDE (40) to determine indel spectra and to estimate the changes in TIF upon drug treatment. AT9283, PCI-24781 and Tubastatin A increased TIF to similar extents as observed in the screen (**Figure 2D, S2D**), while Resveratrol did not cause an increase in TIF in any locus. These results validate the identified TIF-boosting drugs (except Resveratrol) and show that these drugs, especially AT9283, are potent in increasing gene editing efficiencies in heterochromatic loci.

We then looked at the strongest TIF-decreasing drugs. We observed that decreases in TIF generally correlate with reduced viable cell counts (Pearson’s R = 0.63, *p* < 2.2 × 10^-16^) (**Figure S1B, S2A**). To limit this potentially confounding factor we increased the viable cell count threshold for compounds that showed decreased TIF from 25% to 70% (Pearson’s R = −0.0062, *p* = 0.9) (**Figure S2B**). Among the drugs that pass this more stringent filter, we found mainly Aurora Kinase inhibitors (n = 5), all targeting Aurora Kinase A. Aurora Kinase A has a role in cell cycle regulation as well as DSB repair (41,42). Both these mechanisms could explain the observed phenotype. Besides Aurora Kinase A inhibitors, two epigenetic reader domain (ERD) inhibitors, more specifically BRD4 inhibitors, caused the strongest global decrease in TIF (**Figure 2C, S2B**). BRD4 inhibitor-mediated decreases in TIF might be attributed to a global decrease in transcription output, which may also result in reduced expression levels of Cas9 (e.g.,43); or effects on histone eviction (44); or a role of BRD4 in DNA repair (45,46). Drugs that reduce TIF are not of practical use for improving Cas9 editing but might provide insight into the determinants of TIF.

We then investigated the local effects of the drugs on the IPRs, which revealed several notable patterns. First, we found that the high efficiency of this sgRNA was only slightly altered in euchromatin compared to heterochromatin (**Figure 2E**). This phenotype is most likely caused by saturation of TIF at these loci. Second, we observed that many drugs showed complex and IPR-specific changes in TIF, suggesting that drugs interact with the chromatin context in a specific manner. Interestingly, drugs that target the same protein group often showed similar IPR-specific patterns and clustered together, which is demonstrating the chromatin-dependencies of these target proteins. Moreover, several drugs showed concentration-dependent effects, with the higher concentrations often leading to a stronger decrease in TIF along with lower cell count (e.g., I-BET-151). These changes can also be chromatin-dependent; e.g., the DNA methyltransferase inhibitor Decitabine increases TIF globally at 100 nM but has a mixed effect at 1 µM where it decreases the TIF in triple heterochromatin and increases it in euchromatin (**Figure 2E**). This mixed effect was also present for three HDAC inhibitors: Apicidin, Vorinostat and Droxinostat. Together, this shows that the effect of chromatin-modifying drugs is concentration- and chromatin-dependent.

### TIF-modulating drugs display a diversity of chromatin context-dependencies

Visual inspection of **Figure 2E** suggested that many of the tested drugs have quantitatively different effects across the 19 IPRs. We applied a statistical framework to systematically identify such chromatin context-dependencies of the drug effects on TIF, taking advantage of the fact that for each IPR location the local levels of 25 different chromatin features are known (**Figure 1A**). Our aim was to first identify the drugs with statistically significant dependency on the overall chromatin composition, and then to identify the individual chromatin features that may explain these dependencies (see Methods). For the first step we used principal component regression. This approach was taken because many chromatin features co-variate, necessitating a dimension reduction approach. This yielded 49 drugs and 66 drug/concentration combinations with significant chromatin context-dependencies. For each of these drug/concentration combinations we then fitted the log_2_-fold TIF changes to each of the 25 chromatin features by standard linear modelling. The slopes of these linear fits provide a measure of the degree to which each chromatin feature can explain the magnitude of the drug effect (**Figure 3A-C**). We focused on chromatin features with positive slopes, which are candidates to boost the TIF-enhancing effects of a drug (**Figure S3A, S3B**). A total of 35 drugs (41 drug/concentration combinations) exhibited such positive slopes (**Figure 3D**).

**Figure 3.**
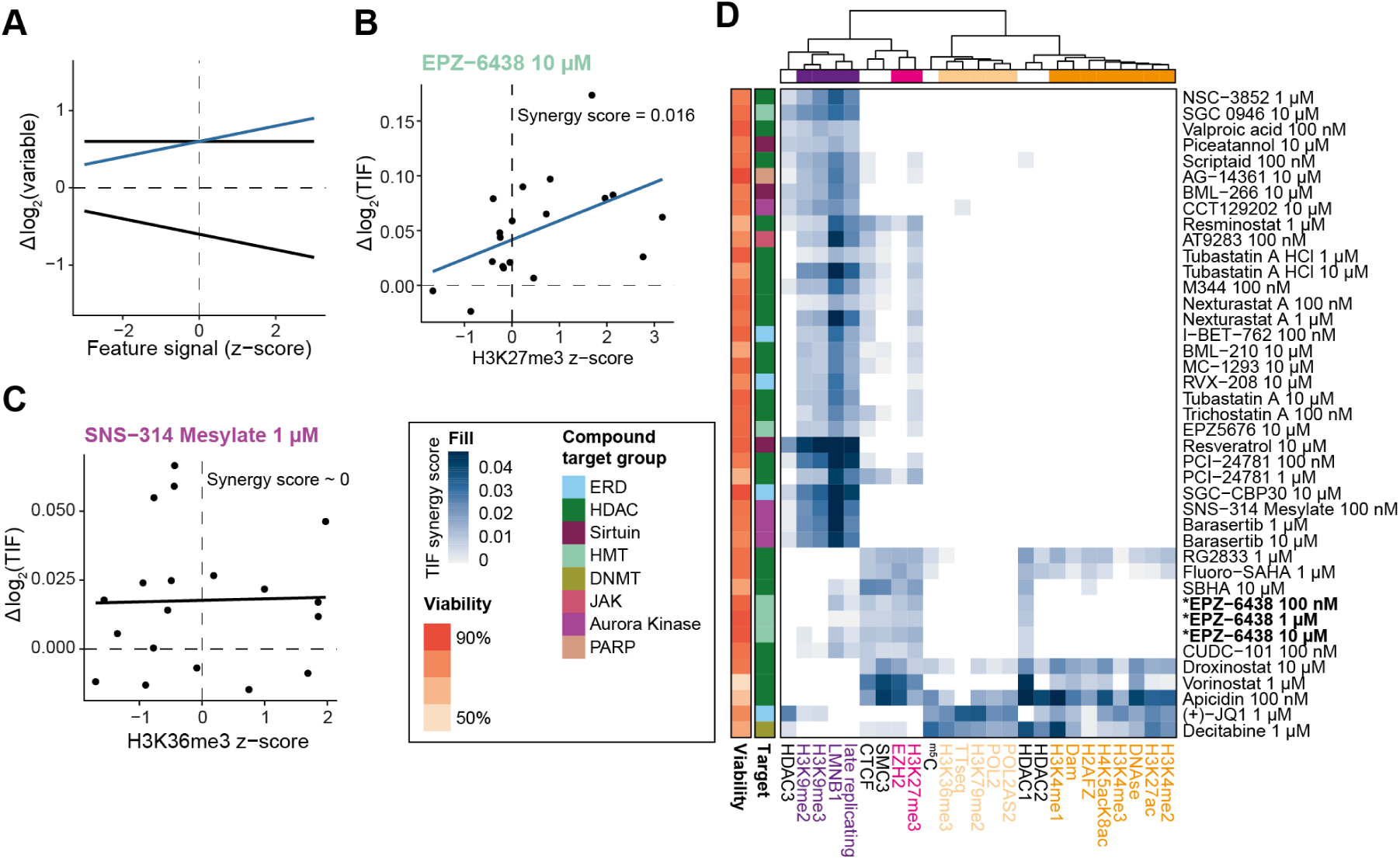
Chromatin context-dependent (CCD) effects for drugs affecting TIF. (A) Example of linear regressions of drug effect-chromatin feature synergies. Highlighted in blue is a positive synergy, no or a negative synergy is shown in black. (B) Example of a positive synergy between the Δlog2(TIF) after EPZ-6438 treatment and the H3K27me3 chromatin feature. The indicated synergy score represents the slope of the linear model. (C) Example of a neutral effect; the treatment with SNS-314 Mesylate has no synergy with H3K36me3. (D) Heatmap of all the significant positive CCDs for the drugs.

A striking and readily interpretable example of such a positive chromatin context-dependency (CCD) is EPZ-6438, an inhibitor of the H3K27 methyltransferase EZH2 (**Figure 3B**). Although this drug enhances TIF at all 19 IPRs (**Figure 2E**), it preferentially enhances TIF in regions rich in H3K27me3 and EZH2 binding. This is the case for all three tested concentrations (**Figure 3D**). EPZ-6438 also shows a synergy with late-replicating regions, lamina-associated regions and regions bound by CTCF or the cohesion subunit SMC3. CTCF and SMC3 have been reported to partially overlap with H3K27me3-marked domains (47,48) and indeed the two IPRs in clone #5 with high CTCF and SMC3 scores overlap with H3K27me3 (**Figure 1A**). Therefore, this CCD should be interpreted with caution. We suggest that depletion of H3K27me3 by EPZ-6438 facilitates access of Cas9 by locally relaxing the heterochromatic state.

Besides the EZH2 inhibitor EPZ-6438, various other drugs showed positive synergies with H3K27me3 (**Figure 3D**). Interestingly, these drugs are primarily HDAC inhibitors (**Figure 4A)**, which may be related to the mutual exclusivity of H3K27me3 and H3K27ac (49). Moreover, HDAC inhibitors exhibited additional CCDs and could be generally divided into two groups based on their CCDs. Some HDAC inhibitors (e.g., PCI-21781, Tubastatin A, Valproic acid) showed prominent synergy with features that mark triple heterochromatin. Others (e.g., Apicidin, Droxinostat, Vorinostat) enhanced TIF in both euchromatic regions and H3K27me3 regions (**Figure 4B**). Since HDAC inhibitors target HDAC proteins with varying specificities, we wondered whether the observed chromatin dependencies may be explained by HDAC protein specificities. However, a survey of previously characterized HDAC specificities (38) (**Figure 4B**), did not uncover any obvious link between the specificity of the HDAC inhibitor and the context-dependencies of the TIF effects (**Figure 4B**). As may be expected, several HDAC inhibitors are more effective in regions bound by HDAC1, 2 or 3 (**Figure 4B**), but this is not true for all HDAC inhibitors. However, most HDAC inhibitors target multiple HDACs, including ones for which no genome-wide binding maps are available.

**Figure 4.**
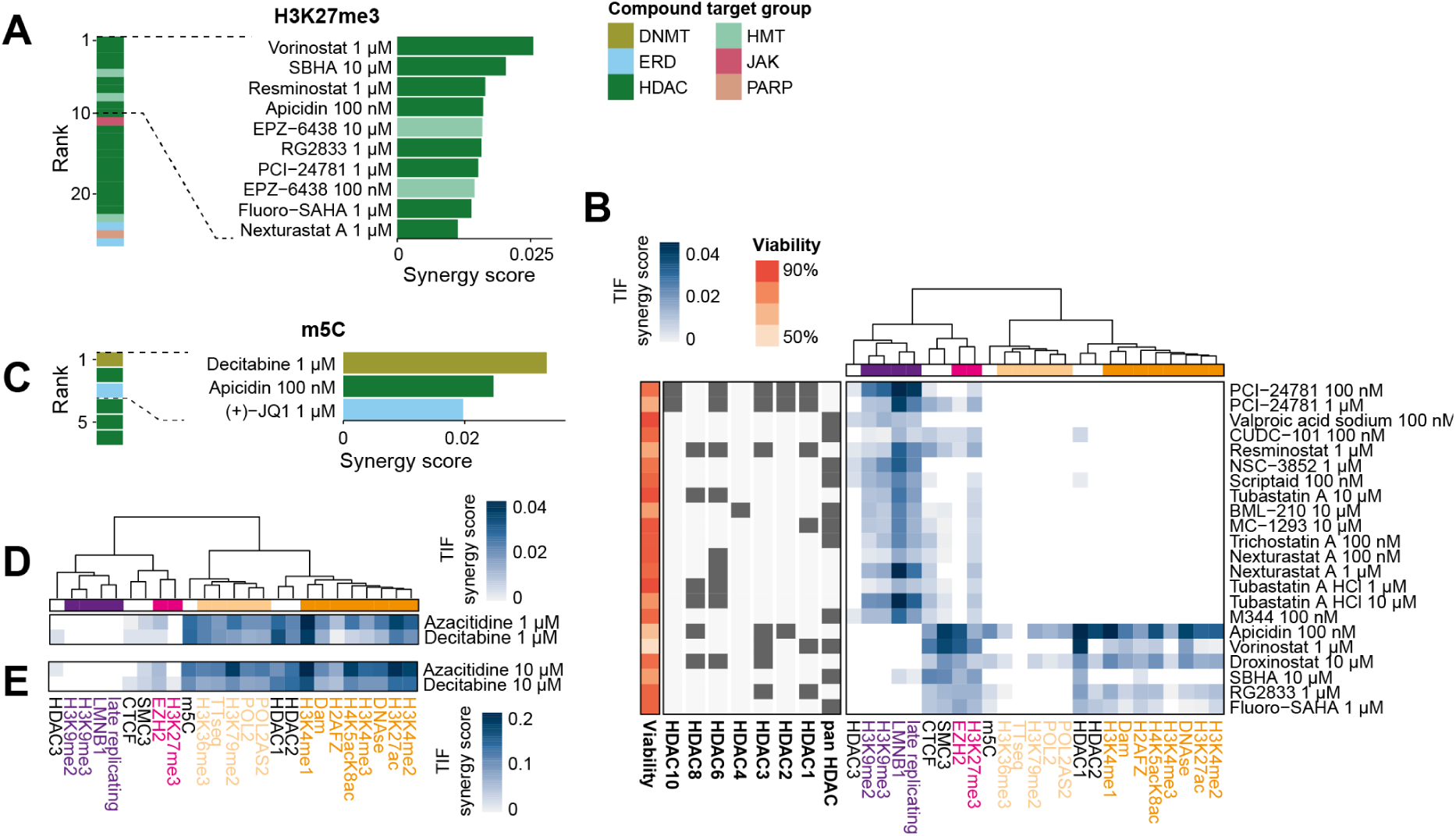
Chromatin context-dependency analysis reveals drugs boosting TIF in specific contexts. (A) Highest positive synergies with H3K27me3, highlighting the top ten drugs with significant interactions. Target categories of the drugs are indicated by color. (B) Heatmap of all the significant synergies of HDAC inhibitors with the 23 chromatin features. Row annotation includes relative viable cell count and the HDAC target for each drug as reported by the manufacturer (SelleckChem). Pan HDAC are general HDAC inhibitors without specific targets, additional HDAC annotations indicate the main targets. (C) Same as in A but for ^m5^C, highlighting the three top hits. (D, E) All chromatin synergy scores for the two DNMT inhibitors at 1 or 10 µM with significant CCDs.

Interestingly, for genomic regions marked by 5-methylcytosine (^m5^C) the drug with the strongest synergy was the DNA methyltransferase inhibitor Decitabine at a concentration of 1 µM (**Figure 4C, Figure S4A**). It is possible that ^m5^C impedes interactions of Cas9 with its target sequence, and that Decitabine alleviates this effect by removal of ^m5^C. Indeed, some studies found reduced Cas9 editing efficiency in ^m5^C marked sequences (5,13,16) but other reports concluded that ^m5^C does not impede Cas9 binding and cleavage (50 Fujita, 2016, 27465215). We note that the preferential effect of Decitabine and another DMNT inhibitor Azacitidine is not restricted to ^m5^C-marked regions, but also correlates with multiple euchromatic features (**Figure 3D, Figure 4D, E**). We observed a similar TIF-enhancing effect across euchromatic features for (+)-JQ1, a BET bromodomain inhibitor (**Figure 3D)**. However, this preference was not mirrored by two other BET inhibitors, RVX-208 and I-BET-762, which both enhanced TIF preferentially in triple heterochromatin, albeit at distinct concentrations (**Figure 3D)**. Finally, several Aurora Kinase B inhibitors showed strong TIF-enhancing effects in triple heterochromatin (**Figure 3D)**. This may be related to the regulatory roles of Aurora Kinase B in heterochromatin (51).

Together, these results reveal striking differences in the activities of epigenetic drugs in distinct chromatin contexts. Thus, to improve Cas9 genome editing, a drug should be selected depending on the chromatin context of the target sequence.

### Effects of epigenetic drugs on MMEJ:NHEJ ratio

We then analyzed how drugs affect the DNA repair pathway balance. Strikingly, only four drugs significantly increased the MMEJ:NHEJ ratio globally in addition to the NHEJ inhibitor NU7441, which served as positive control in this analysis (**Figure 5A, S5A**). These included two formulations of the PARP-1 inhibitor PJ34 as well as the JAK and Aurora Kinase inhibitor AT9283 and the Aurora Kinase B inhibitor Barasertib.

**Figure 5.**
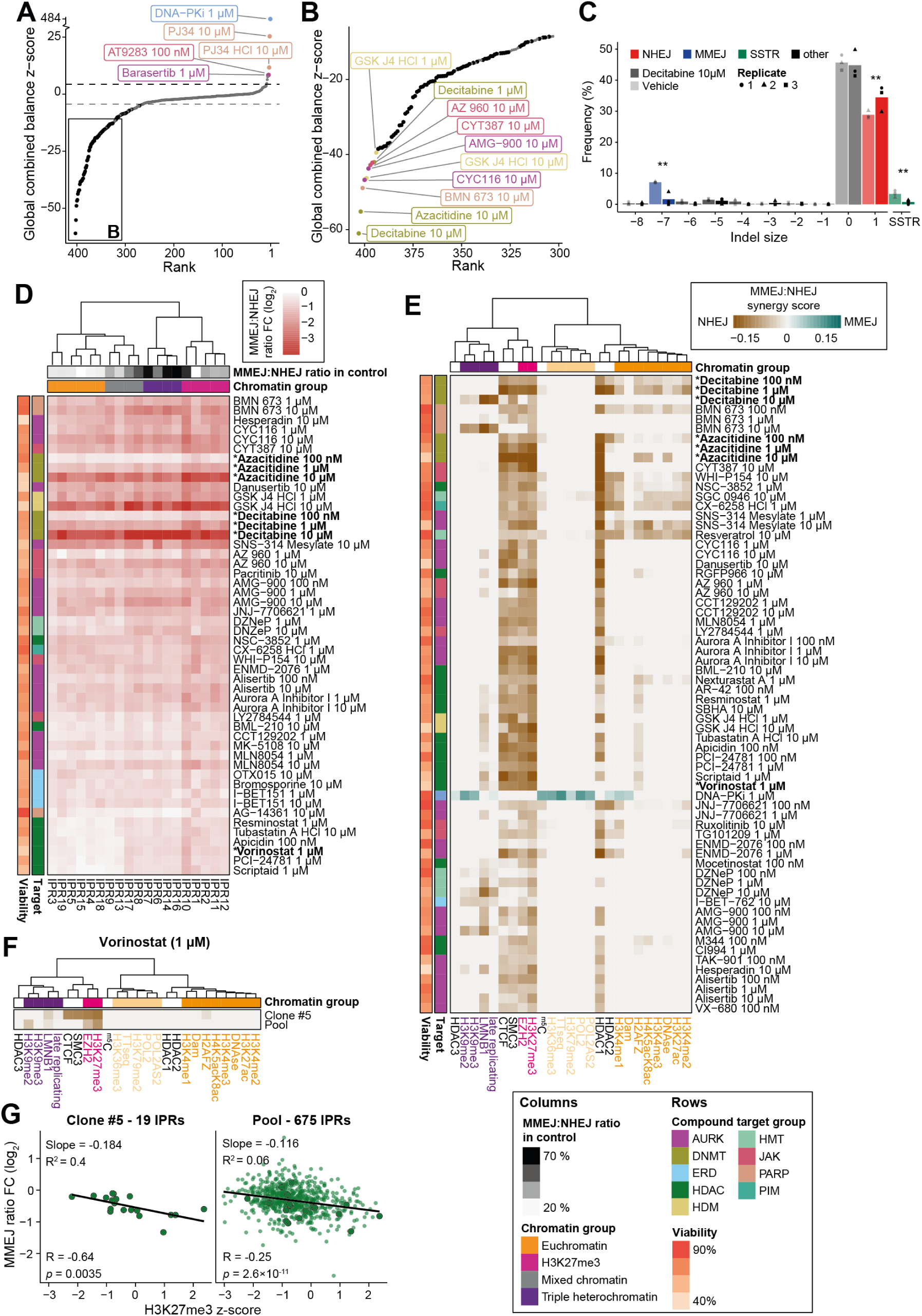
Various drugs are potent inhibitors of MMEJ. (A) Rank dot-plot of the drugs ranked on their global MMEJ:NHEJ ratio z-score. Black dots are samples with a significant global effect in both screens, grey dots are non-significant or only significant in one screen. Top 5 hits increasing the MMEJ:NHEJ ratio globally are highlighted. Dashed lines indicate significance threshold of *padj* < 0.01. (B) Zoom in of A, the bottom 10 hits are highlighted (C) TIDE results of cells treated with Decitabine. The shapes represent the different replicates, the mean is indicated by the bar. Asterisks denote p-values based on Benjamini-Hochberg adjusted T-tests, * = *padj* < 0.05, ** = *padj* < 0.01. (D) Heatmap of the top 60 drugs altering the MMEJ:NHEJ ratio. (E) Heatmap showing all significant chromatin interactions with the MMEJ:NHEJ ratio fold change (FC). (F) Heatmap of the chromatin interactions with the MMEJ:NHEJ ratio FC upon Vorinostat treatment in clone #5 and the IPR pool. (G) Scatter plot demonstrating the synergy between the Vorinostat-induced MMEJ:NHEJ ratio change and the H3K27me3 z-score in the single IPR clone and the complete IPR pool. Transparent dots show IPRs of the pool, dots with black stroke show IPRs of the clone. Linear fit is shown as black line with slope and R^2^ above. Denoted below the fit are Pearson’s R and correlation significance.

In contrast, 95 drugs (102 drug/concentrations) caused a globally decreased MMEJ:NHEJ ratio (**Figure 5B**). Mirin, an inhibitor of Mre11, was used as positive control for the decrease of MMEJ but had only little effect and was not reproduced in all replicates. The large number of drugs decreasing the MMEJ:NHEJ ratio suggests that the NHEJ repair pathway is much more robust than the MMEJ pathway. Because the activity of MMEJ is known to be regulated during the cell cycle and requires passage through mitosis (52,53), we investigated the cell cycle profile for 25 drug/concentration combinations with a significant effect on the MMEJ:NHEJ ratio. While most of the Aurora Kinase inhibitors induced a considerable G2 arrest, most drugs targeting other proteins did not substantially alter cell cycle profiles (**Figure S5B**). It is possible that the MMEJ is more easily perturbed because it is a more complex repair pathway that relies on the resection machinery (21,54–56).

Among the drugs with the strongest reduction in the MMEJ:NHEJ ratio were the DNMT inhibitors Decitabine and Azacytidine (**Figure 5B**). These drugs almost completely blocked MMEJ in all IPRs. Decitabine and Azacytidine are cytidine analogues that are incorporated into DNA upon replication and then covalently trap DNMTs on the DNA (57). We wondered whether this may interfere with resection or DNA synthesis, which are key steps of MMEJ. We therefore investigated whether Decitabine could inhibit another resection-dependent pathway, single-strand template repair (SSTR). We tested this by conducting Cas9 editing of the *LBR* gene in the presence of a single-stranded oligonucleotide template that directs a specific 2 bp insertion (15). We used TIDE (40) to quantify the resulting indel pattern. According to this assay the activities of both MMEJ and SSTR, but not of NHEJ, were significantly reduced in Decitabine-treated cells (**Figure 5C**). This result suggests that Decitabine selectively blocks resection-mediated repair, although it does not identify the affected step of this process.

Visual inspection of the changes of MMEJ:NHEJ balance per IPR (**Figure 5D**) suggested that the changes in MMEJ:NHEJ balance triggered by various drugs may not be uniform across the different chromatin contexts. We therefore subjected the changes in MMEJ:NHEJ ratio to a similar analysis of chromatin context-dependence as described above for TIF changes, except that we now searched for chromatin features that correlate with either increased or decreased MMEJ:NHEJ ratios (**Figure 5E**). This yielded 43 drugs (65 drug/concentration combinations) with significant chromatin context-dependencies in addition to DNA-PK inhibition with NU7441. Strikingly, all these drugs, except for NU7441, preferentially reduce the MMEJ:NHEJ ratio in regions marked by H3K27me3. Many of these are also more effective in the IPRs with higher levels of EZH2, CTCF and SMC3, and HDAC1. This suggests that the MMEJ:NHEJ pathway balance in these regions is easily shifted by a broad variety of perturbations. Among the drugs synergizing with H3K27me3, Aurora Kinase inhibitors are enriched (Fisher’s exact test *p_adj._* = 6.93 × 10^-4^), however their general effect on DNA repair and the cell cycle might be a confounding factor. Additionally, many HDAC inhibitors showed the strongest synergistic effect with H3K27me3 (**Figure S5C**). Interestingly, HDAC inhibitors typically reduced the MMEJ:NHEJ ratio less efficiently in euchromatic regions compared to other effective drugs (**Figure S5D**). Again, this H3K27me3-specific phenotype of HDAC inhibitors might be related to the mutual exclusivity of H3K27me3 and H3K27ac (49). Besides H3K27me3 interactions, some drugs showed an additional preference for euchromatic regions rich in regulatory elements, but not in transcribed regions. These drugs included Decitabine at lower concentrations (100 nM and 1 µM) and Azacitidine, but also a diversity of other drugs with no obvious commonality. Six drug/concentration combinations preferentially reduced the MMEJ:NHEJ ratio in triple heterochromatin (**Figure 5E**), but again without an obvious pattern in their target specificities.

Although the 19 IPRs in clone #5 covered all major chromatin features, we considered the possibility that they were not fully representative of the genome-wide diversity of chromatin states. This could confound the interpretation of our data. To test whether the results of the 19 IPRs could be generalized to other genomic regions, we employed a previously established pool of K562 cells with 675 randomly integrated IPRs of which the local chromatin composition is known (15). We treated this cell pool with 1 µM Vorinostat, an HDAC inhibitor with local MMEJ:NHEJ ratio changes (**Figure 5E**), and determined the MMEJ:NHEJ ratio in each IPR. Interestingly, we found a correlation between the H3K27me3 and EZH2 scores and the Vorinostat-induced changes in MMEJ:NHEJ ratios as observed with clone #5 (**Figure 5F, 5G**). The correlations with SMC3 and CTCF, however, could not be observed in the pool. This demonstrates that the synergy of SMC3/CTCF observed in clone #5 is likely caused by co-occurrence with H3K27me3 and that the true causal feature is H3K27me3. Based on these results we conclude that DSBs in H3K27me3-rich chromatin are susceptible to changes in the MMEJ:NHEJ pathway balance as previously reported (15).

## DISCUSSION

The screen presented here provides new information on how drugs alter Cas9 editing efficiency and DNA repair outcomes in the light of the chromatin environment. Importantly, besides the known HDAC inhibitors, we find many novel drugs that can be used to improve gene editing efficiency. Additionally, we highlight that the improvement of Cas9 efficiency depends on the inhibitor and the type of chromatin. The performed chromatin context-dependency (CCD) analysis (56) demonstrates in detail how the various drugs interact with chromatin and how that changes the repair pathway balance and Cas9 efficiency. For example, while the two HDAC inhibitors PCI-24781 and Apicidin both on average increase TIF, only PCI-24781 efficiently does so in triple heterochromatin. Apicidin, on the other hand, only increases TIF in H3K27me3-marked chromatin and even decreases TIF in triple heterochromatin. These distinctions may aid users in choosing drugs that work best for their CRISPR target region. It is likely that drugs that increase TIF in specific regions are preferable to use over drugs that increase TIF globally, as the latter could potentially increase off-target editing.

The effect sizes that we observed were mostly modest. This is at least in part since the sgRNA we use in this study is extremely efficient, leading to near saturation of the TIFs in accessible regions (15). A less efficient sgRNA targeting the reporter would presumably have yielded a higher dynamic range, and therefore the effects of the drugs on TIF may be more pronounced with less efficient sgRNAs.

There are some observations that would require further exploration. Aurora Kinase inhibitors were found to increase and decrease both the TIF and the MMEJ:NHEJ balance (**Figures 2B-C & 5A-B**). Strikingly, Aurora Kinase B inhibitors such as Hesperadin and Barasertib were found to boost the TIF and shift the balance towards MMEJ respectively, while Aurora Kinase A inhibitors had the opposite effect. Whether these are direct or indirect effects is unclear, but both kinases are involved in DNA repair and cell cycle progression (42). This result could be confounded by the relative selectivity of these drugs for Aurora Kinase A and/or Aurora Kinase B as many drugs target both kinases at different concentrations. We also found that many Aurora Kinase inhibitors had a stronger effect on the repair pathway balance in the presence of H3K27me3. It is possible that Aurora Kinases are involved in the repair signaling of DSBs occurring in H3K27me3 regions.

This screen also helped to identify strong inhibitors of MMEJ and identify interactions with chromatin features for a multitude of drugs. We show that many drugs perturb MMEJ and that they do so preferentially in H3K27me3-marked regions. We also demonstrate that Decitabine is a strong inhibitor of two resection-based repair pathways.

While further validations of many of these drugs are needed to obtain a better control and understanding of Cas9 editing and DNA repair, this screen should provide a valuable resource for future DNA repair studies and further improvements of gene editing technologies.

## MATERIALS & METHODS

### Generation of cell line and cell culture

All experiments were performed on the earlier established K562#17, which is a clonal cell line of the female K562 cells (ATCC) stably expressing DD-Cas9 (20,39). In this study, we used a heterogenous DSB-TRIP cell-line, harboring thousands of DSB reporters, and for the screening we selected a clone (clone #5) carrying 19 reporters (15,58). Both the clone and cell pools were cultured in tissue culture flasks in RPMI 1640 (GIBCO) supplemented with 10% fetal bovine serum (FBS, HyClone) and 1% penicillin/streptomycin. Cells were kept at 37°C in a humidified atmosphere containing 5% CO_2_. For both screens, three separate cryovials of clone 5 cells were cultured separately for two weeks, one for each replicate. The TRIP cell pools were kept in culture for less than two weeks to avoid reporter drifting (58).

### sgRNA transfections

To induce DSBs at the 19 sgRNA target sites, per replicate, 30 × 10^6^ clone 5 cells were transfected with the sgRNA plasmid (five nucleofections of 6 × 10^6^ cells), and 6 × 10^6^ clone 5 cells for the control nucleofection (plasmid carrying GFP). For transfections, 6 × 10^6^ clone 5 cells were resuspended in 100 μL transfection buffer (100 mM KH2PO4, 15 mM NaHCO3, 12 mM MgCl2, 8 mM ATP, 2 mM glucose (pH 7.4)) (59). After the addition of 6.0 μg plasmid DNA expressing either LBR2 sgRNA or GFP, the cells were electroporated in an Amaxa 2D Nucleofector using program T-016. Subsequently, the electroporated cells were plated in 10-cm dishes and incubated overnight for the cells to recover. Nucleofection efficiency was estimated by flow cytometry on the GFP transfected samples prior to the screening.

### Addition of the epigenetic screening library

10^4^ cells per were dispensed in each well of a 96-well plate in a total volume of 80 μL using a Multidrop Combi fluid dispenser (ThermoFisher Scientific). For the drug addition, we combined two 96-well epigenetic drug screening libraries from two commercially available libraries from Selleck Chemicals and Enzo Life Sciences (for list of drugs see **Table S1**). The drugs had stock solutions of 5 mM, 1mM and 500 μM and were diluted in a step wise process in complete medium with 500 nM final concentration Shield-1 for Cas9 stabilization (Aobious cat. no. AOB1848) (39), and then added to the sgRNA transfected cells, yielding final drug concentrations of 10 μM, 1 μM, and 100 nM. All steps were performed using a Microlab STAR liquid handling workstation (Hamilton). Four different control drugs were added to the outer columns of the 96-wells using a HP D300 Digital Dispenser. DNA-PKcs inhibitor NU7441 (Cayman; diluted 1:1000 from 1 mM stock in dimethylsulfoxide [DMSO], final concentration 1 μM) was used as a positive control for NHEJ inhibition, MRE-11 inhibitor Mirin (diluted 1:1000 from 12.5 mM stock in DMSO, final concentration 12.5 μM) was used as a positive control for MMEJ inhibition, DMSO was used as a negative control, and Phenylarsine Oxide (PAO; 10 μM) was used as a positive toxicity control.

### Viable cell count readout

The viable cell count was measured 72 hours after the drug addition with a resazurin assay. 20 μL of 5x resazurin was added to the 96-wells plate with the Multidrop Combi fluid dispenser (ThermoFisher Scientific). Plates were shaken and incubated for 3h at cell culture conditions before measuring the fluorescence (560Ex/590Em) on an EnVision Multilabel Plate Reader (Perkin Elmer). The lowest fluorescence from the PAO killing control in each replicate was subtracted from all the samples within the sample replicate to standardize the raw values. The viable cell counts per sample were then normalized on the average fluorescence of the control samples (DMSO treated) within each of the six replicates. Cell count viability filters are further explained below in the computational analysis section.

### High-throughput sequencing

After measuring the viable cell count, the cells were processed for PCR. The cells were transferred to 96-well plates suitable for PCR and subsequently centrifuged at 300 g for 5 min. Then, the supernatant was removed prior to the addition of 20 μL DirectPCR Lysis Reagent Cell (Viagen, cat. no. 301-C) supplied with 1:100 10 mg/mL Proteinase K (Bioline, cat. no. BIO-37084). The cells were lysed by incubating for 3h at 55°C and the Proteinase K was inactivated for 10 min at 95°C. PCRs were performed in two steps: PCR1 with 10 μL of the crude lysates, 25 μL 2 x MyTaq HS Red Mix (Bioline, cat. no. BIO-25048), and plate-specific Illumina PCR indexed primers (TAC0012 and TAC0007, final concentration 1 μM; **Table S2**). PCR2 was performed using 15 μL 2 x MyTaq Red Mix (Bioline, cat. no. BIO-25044) and 5 μL of each PCR1 product with well-specific Illumina PCR indexed primers (TAC0159 & TAC0009, 1 μM; **Table S2**). Each sample was generated with a unique combination of a plate-specific primer (TAC0007) and well-specific primer (TAC0159). PCR1 was carried out using initial 5 cycles of 1 min at 95°C, 15 s at 95 °C, 15 s at 55 °C and 15 s at 72°C; followed by 10 cycles of 15 s at 95 °C, 15 s at 70 °C and 15 s at 72°C. PCR2 was performed using one of the initial cycles; followed by 14 of the subsequent cycles. 10 μL of all wells of three different 96-wells plates were loaded on a 1% agarose gel to check for variability between individual wells. Subsequently, all wells were pooled equally for each well using 5 μL from each reaction. Primer dimers were removed by CleanPCR bead purification (CleanNA, cat. no. CPCR-0050) using a 0.8:1 bead:sample ratio. The samples were then eluted in 50 μL and further purified by loading it onto a 2% agarose gel. The PCR product was cut from gel to remove any other undesired products and cleaned with the PCR Isolate II PCR and Gel Kit (Bioline, cat. no. BIO-52060). The isolated samples were sequenced by Illumina MiSeq for screen 1 (14.3 million reads) and by NextSeq Mid for screen 2 (88.4 million reads) which allowed for an average of 477 reads per IPR per sample in screen 1 and 2,896 reads in screen 2.

### Flow cytometry to assess cell cycle arrest

To generate cell cycle profiles upon drug addition, the DNA content was measured using flow cytometry. The cells were plated in 96-well plates and damage was induced as described before. Candidate drugs were selected from the epigenetic drug library and added as previously mentioned. 72 hrs after the Cas9 activation and drug addition, the cells were centrifuged at 300 g for 5 min. Subsequently, the supernatant was removed followed by resuspension in 140 μL Nicoletti buffer (0.1% sodium citrate (pH 7.4), 0.1% Triton X-100, 50 μg/mL propidium iodide (PI)) supplied with 1:100 10 μg/mL RNase A (ThermoFisher Scientific). The DNA content of the samples in the 96-wells plates was measured on the Attune NxT Acoustic flow cytometer (ThermoFisher Scientific) using a 405 nm laser for excitation. Flow cytometry data and cell cycle fractions were analyzed and calculated using FlowJo v10.

### TIDE experiments to estimate Decitabine-induced changes in indel ratios

To examine DSB repair pathway choice upon treatment with Decitabine, we induced DSBs using our sgRNA in clone #5 cells and then added DMSO or 10 μM of Decitabine. To assess SSTR, we transfected together with the sgRNA a single-stranded oligodeoxynucleotide (ssODN) that served as a template for SSTR. This template was previously characterized and typically leads to a +2 insertion at the sgRNA target site (15). As a negative control, the cells were transfected with a GFP-expressing plasmid. The Cas9 was activated by the addition of Shield-1. After 72h, DNA was isolated using 20 μL of DirectPCR Lysis Reagent Cell (Viagen) supplemented with 1:100 10 mg/mL Proteinase K (Bioline). TIDE was performed as described before (40). The sgRNA target site was amplified by PCR (using primers TAC0017 & TAC0018), purified, and subjected to Sanger sequencing (with TAC0018). Sequencing results of the GFP transfected cells and the sgRNA transfected cells were then analyzed using TIDE.

### TIDE experiments to estimate TIF in additional heterochromatic loci

To assess drug-induced increases in TIF in other genomic loci than the one used in the screen, we selected four additional sgRNAs targeting heterochromatic loci (**Table S3**). Per sgRNA, 10^6^ K562 clone #5 cells were transfected using 2 μg of sgRNA-expressing plasmid. After 12 hours, 2×10^4^ cells were plated in 96-wells and Shield-1 and drugs (DMSO, PCI-24781 (100 nM), AT9283 (100 nM), Resveratrol (10 μM), Tubastatin A (10 μM)) were added. 72h later, DNA was isolated using 20 μL of DirectPCR Lysis Reagent Cell (Viagen) and sgRNA target sites were amplified for TIDE analysis as described above (using primers SM8+9 (LAD2), SM18+19 (LAD7), SM24+25 (LAD10) and TAC0200+201 (TBX5)). Experiments were performed in five replicates on five different days.

### TRIP Pool experiments

The experiments in the cell pools were performed as follows in three biological replicates. The TRIP pools (15) consist of 2 different pools (A and B). 6 × 10^6^ cells from each pool were transfected with 6 μg of LBR sgRNA plasmid as described above and seeded in 10 cm dishes for overnight recovery. Shield-1 and either DMSO (1:1000) or Vorinostat (1 μM final concentration, 1:1000 from 1 mM stock in DMSO) were added to start break induction. The samples were collected 72 hours after the addition of the drugs and genomic DNA (gDNA) was extracted using the ISOLATE II genomic DNA kit (Bioline, BIO-52067). The samples were then quantified using Nanodrop and 200 ng of gDNA was used in the library PCRs that were performed as for the screening above. The resulting libraries were purified and quantified by Qubit before being combined and sequenced by NextSeq MID.

### Analysis of high-throughput sequencing data of the screens

Analysis of the raw demultiplexed sequencing data was done using our previously reported computational pipeline (58). Briefly, indel size counts were generated for each integration of each well. The processed data was then filtered and analyzed in R. Several filtering steps were implemented prior to the analysis. As +1 bp insertions (NHEJ) and −7 bp deletions (MMEJ) were the most important and in general abundant indel reads in our assay, we required each sample to have at least a sum of 30 reads for +1 and −7. Samples with fewer reads were discarded, resulting in 3274 remaining samples (of 3456 total samples), with on average 18.7 IPRs per sample (excluding the PAO samples). The remaining samples were filtered for viable cell counts: samples with a viable cell count score of less than 25% compared to cells treated with DMSO were removed from the analysis to prevent secondary effects. This cutoff was more stringent for analyzing the pathway balance (40%) due to the importance of the cell cycle in the pathway balance choice and even higher for negative effects on the editing efficiency (60%) due to more complex confounding effects (such as transcription, cell cycle). Samples were allowed one replicate to be below the threshold to pass the filter, the others were discarded (65 samples for TIF and 73 samples for the pathway balance were discarded, including the killing control). The TIF and MMEJ:NHEJ ratios were only calculated when the replicate passed the viability cutoffs ({1}, {2}). This made sure we had representative replicates and reliable measurements. They were calculated with the formulas below.

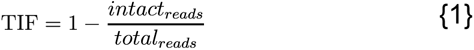

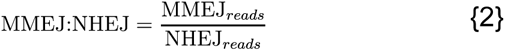

Where *intact_reads_* is the number of reads with 0 indels, *total_reads_* the number of total reads for that sample and barcode, *MMEJ_reads_* the number of 7 bp deletion reads, and *NHEJ_reads_* the number of 1 bp insertion reads.

### Z-score calculation and significance calling

To call significant effect at each IPR for all the samples, we used the 24 control wells per replicate (four per plate and six plates per replicate, from two drug plates and three concentrations) to determine the control distribution per IPR per replicate for the TIF and the MMEJ:NHEJ ratio. We then fitted normal distribution to the control data to determine the mean and standard deviation per IPR per replicate. These were then used to calculate z-scores (*Z_measure_*) for the IPRs in all the samples {3}. The z-scores per IPR, drug, concentration and replicate were calculated as follows for both the TIF and the MMEJ:NHEJ balance.

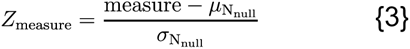

Where measure is either TIF or MMEJ:NHEJ and *N_null_* is a normal distribution of the measure fitted with control treated samples (DMSO, n = 24).

To assure reproducibility between replicates and screens, specifically for the samples with an effect, we set up a two-step reproducibility filter. First, IPRs required an absolute z-score >1.96 (equivalent to *p* < 0.05) in at least one out of two or three replicates or two out of three replicates (on average each IPR was present in 2.9 and 2.87 replicates in screen one and two respectively across all the samples). Second, we calculated the combined z-score of the three replicates in each screen using the Stouffer method (60) {4}. We used this *Z_screen_* only to test whether the replicates had the same effect i.e., the samples required the same sign. We calculated a combined z-score for all the samples as below to obtain the final *Z_combined_* (5). We then transformed this *Z* to a p-value and corrected for multiple testing using the Benjamini-Hochberg correction (61). For the global measurements, we first averaged the values of all the IPRs, and we calculated the z-scores and p-values in the same way. The *Z_screen_* for the screen reproducibility filter was calculated as follows.

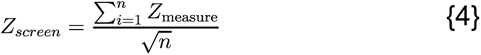

Where n is the number of replicates in that screen with sufficient reads and viable cell count (between one and three).

The final *Z_combined_* was calculated as follows.

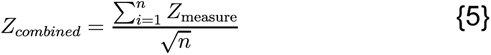

Where n is the number of replicates in total (between two and six).

### Statistical Analyses

The statistical analyses when done are indicated in the text or figure legends. All statistical analyses were corrected for multiple testing as indicated.

### Chromatin context dependency analysis

The chromatin context dependency analysis was performed as described previously (56). We applied a p-value < 0.01 cutoff (after correcting for multiple testing) in this work.

### Chromatin coverage of the validation sgRNAs

The chromatin features for each sgRNA and IPR were processed as reported previously (15) using the coordinates from **Table S3** for the sgRNAs.

## DATA AVAILABILITY

Laboratory notebooks and supplementary data are available on Zenodo (10.5281/zenodo.7866302). Raw sequencing data is available on Sequence Read Archive under BioProject number PRJNA960840. Code and supplementary files used as input for the scripts are available on Zenodo (10.5281/zenodo.7863866).

## ACKNOWLEDGEMENTS

We thank the NKI Genomics, High Throughput Screening, Flow Cytometry, and Research High Performance Computing core facilities for excellent support, and members from our laboratories for inspiring and helpful discussions. We would also like to specifically thank René Medema for his valuable input.

## AUTHOR CONTRIBUTIONS

**Conceptualization:** R.S., M.T, B.v.S., **Data Curation:** R.S., M.T., **Formal Analysis:** R.S., M.T., **Investigation:** R.S., M.T, B.M., **Methodology:** R.S., M.T, B.M., X.V., B.v.S., **Software:** R.S., M.T, B.M., X.V., **Visualization:** R.S., M.T., **Resources:** R.L.B., **Writing – Original Draft Preparation:** R.S., M.T, B.v.S., **Writing – Review & Editing:** R.S., M.T, X.V., B.v.S., **Project Administration:** R.L.B., B.v.S. **Funding Acquisition:** B.v.S. **Supervision:** B.v.S.

## SUPPLEMENTARY FILES

**Figure S1.**
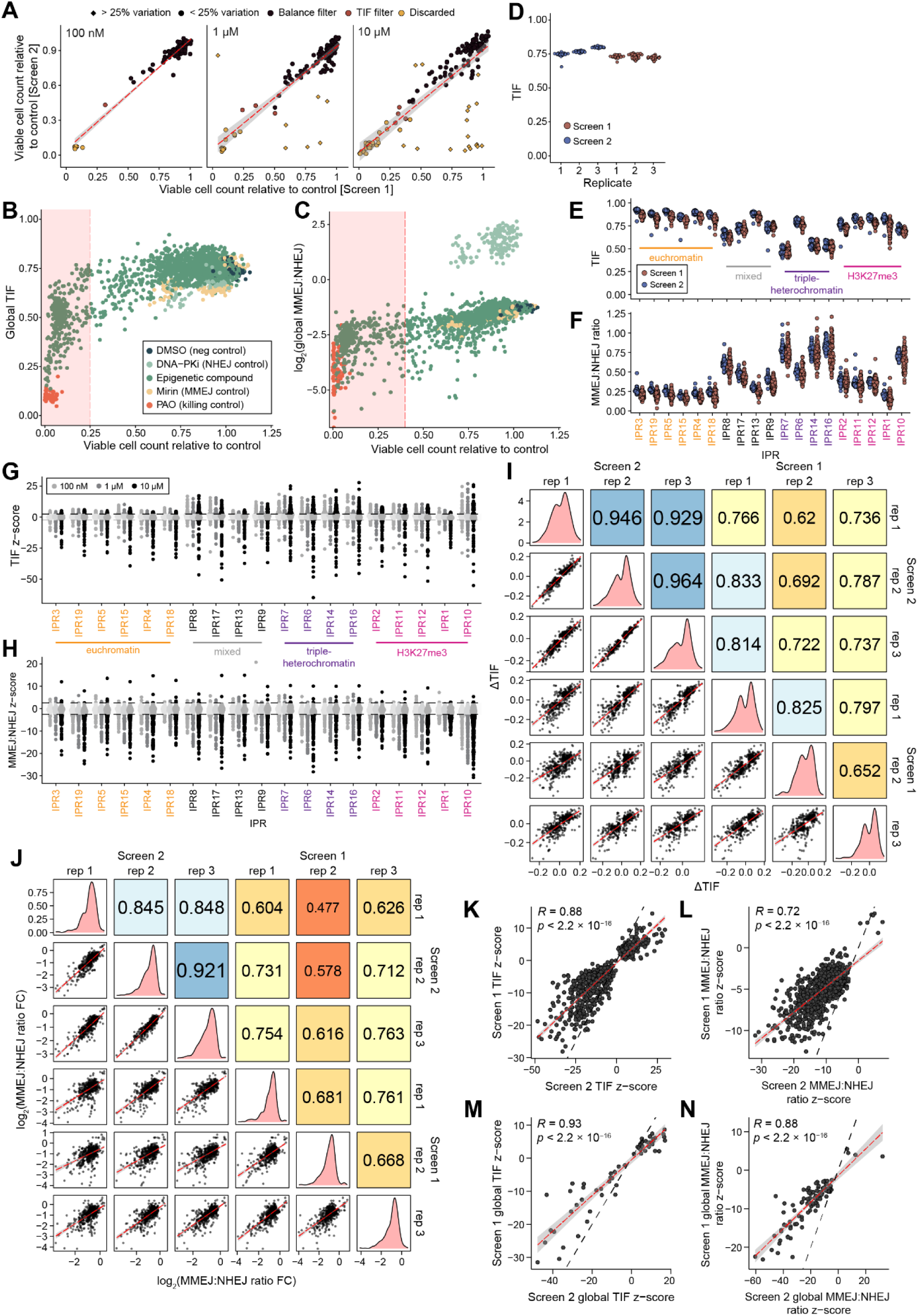
Viable cell count and data reproducibility (related to Figure 1) (A) Scatter plot of relative viable cell counts in the two screens. Black are samples kept for both the pathway balance and the TIF analysis. In orange are samples only kept for the TIF analysis. Yellow samples are discarded. Round samples did not vary more than 25% between replicates, diamond shaped samples varied more and were discarded (yellow). (B) Scatter plot for each separate well (separate replicate and screen) of the relative viable cell count and global TIF. The dots are colored based on drugs (green) and controls (PAO in orange, Mirin in yellow, NU7441 in light green and DMSO in dark green). Samples in the red square are the samples that are discarded with the viable cell count filter. (C) Same as in B but for the global log2(MMEJ:NHEJ). (D) Global TIF per replicate per screen in the vehicle control samples. (E) Average TIF over the three replicates of the vehicle control per IPR for the two screens. The color of the IPR matches the chromatin states based on Figure 1A. (F) Same as in (E) but for the MMEJ:NHEJ ratio. (G) Z-score transformed TIF values. The z-score of the six replicates were combined using the Stouffer’s method (see Material and Methods) (H) Same as in G but for the MMEJ:NHEJ ratio. (I) Correlation between replicates and screens of the ΔTIF for drugs and IPRs that were significant in both screens and present two out of three (or one out of two) replicates each screen. The upper quadrant shows the Pearson’s correlation coefficient. (J) Same as in I but for log2(MMEJ:NHEJ). (K) Scatter plot of all the global TIF z-scores (of the three replicates) between the two screens. Only samples that were significant in both screens are shown. (L) Same as in K but for the MMEJ-ratio z-scores. (M-N) Same as in K-L but for global z-scores (one dot per drug/concentration). For I-M, R is Pearson’s correlation coefficient, the dashed red line and grey shading show the linear regression fit with 95% confidence interval.

**Figure S2.**
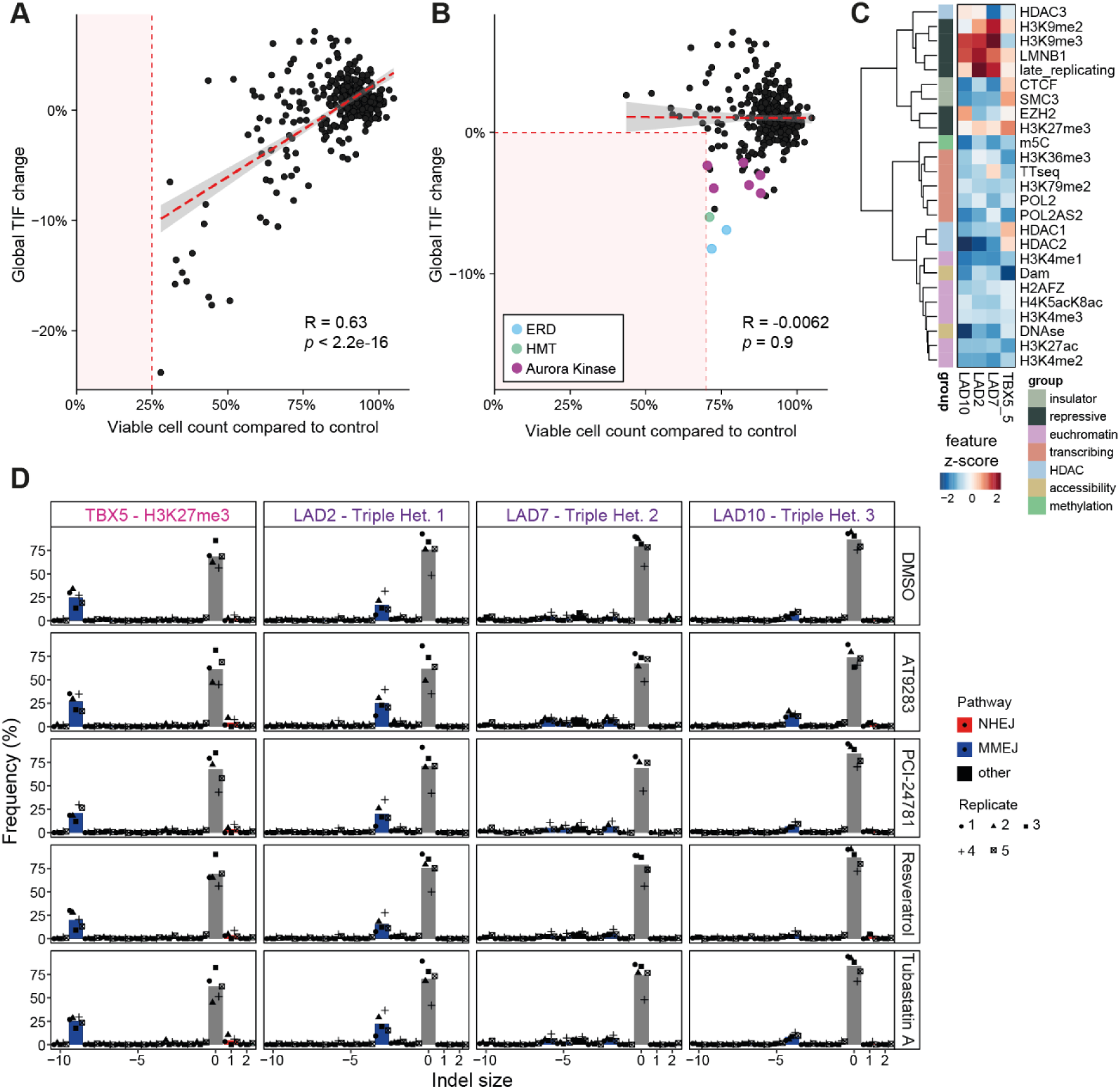
Decrease in TIF could be due to a decrease in viable cell count. (Related to Figure 2) (A) Scatterplot of the relative viable cell count per screen and global TIF change. (B) Same as in A but with a viable cell count cutoff at 70%. The top significant decreasing conditions are highlighted with their target color (Figure 2C). R is Pearson’s correlation coefficient between viable cell count and global ΔTIF for all the samples passing the respective viability filter. (C) Heatmap of the z-scores of the chromatin features in the four tested loci. TBX5 targets a locus with H3K27me3 and the LAD2, LAD7 and LAD10 gRNAs target different LADs. (D) Indel pattern of the four additional heterochromatic loci upon treatment with either DMSO or four of the top hits from the screen. The shapes represent the different replicates, the mean is indicated by the bar. The colors indicate the predominant repair pathways causative for the mutation as predicted by inDelphi (62).

**Figure S3.**
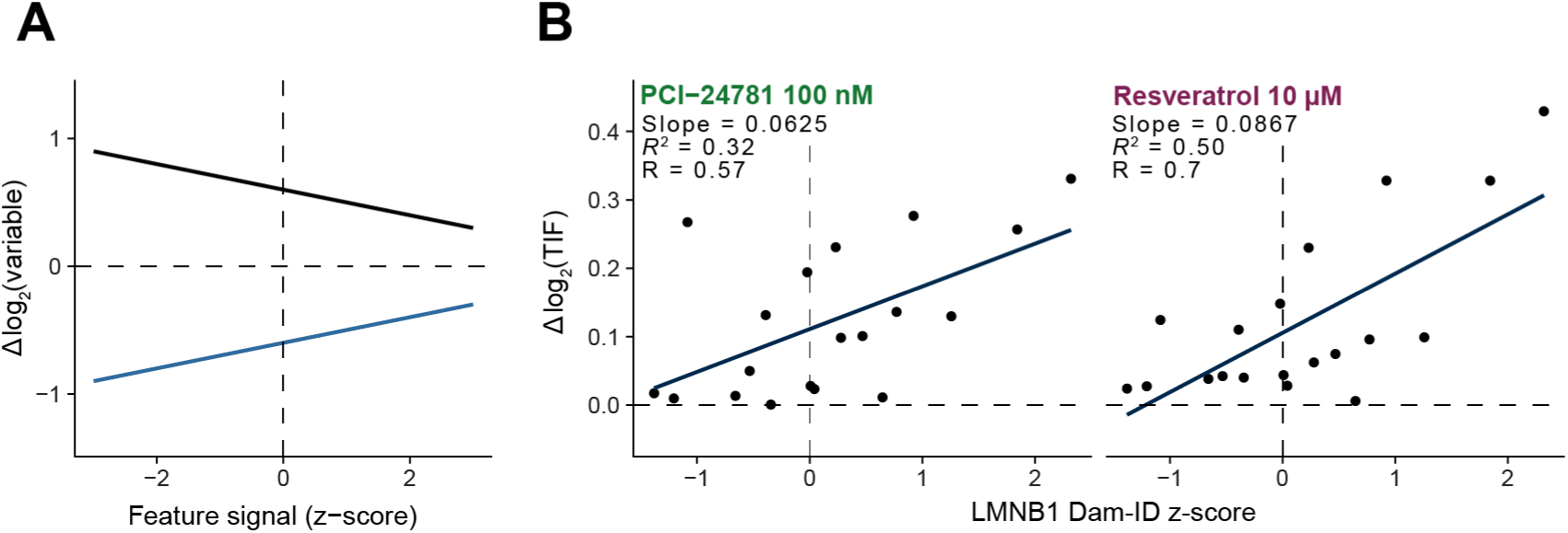
Drugs display specific chromatin context dependencies. (Related to Figure 3) (A) Examples of excluded synergies, with positive effect and a negative (green) slope or vice-versa (blue). (B) Examples of a positive synergy between the effect of the drug and the chromatin feature. Linear model in blue matching Figure 3D, slope and R^2^ of the model and Pearson’s R.

**Figure S4.**
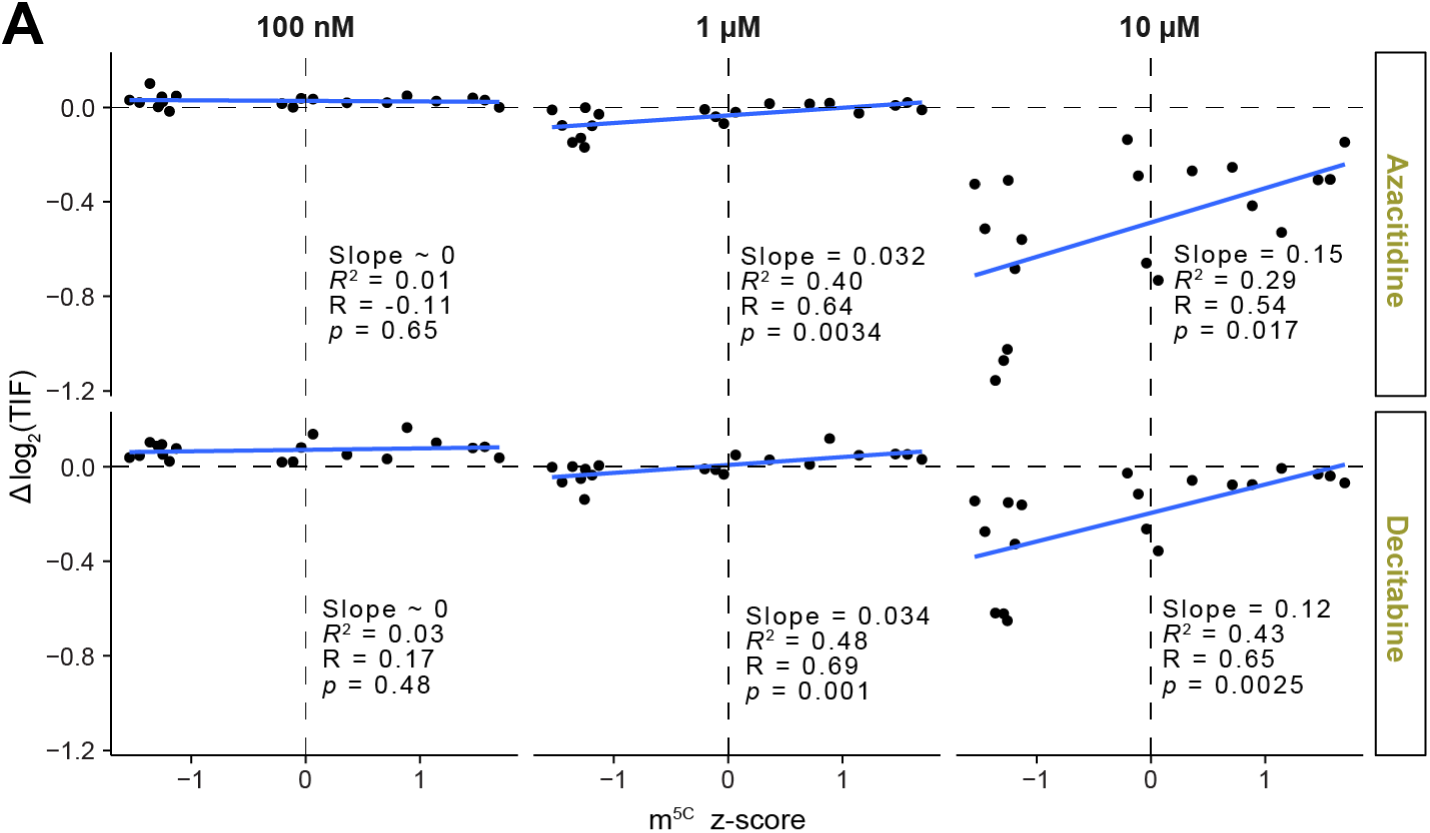
Specific chromatin synergies (related to Figure 4) (A) Scatterplots of the Δlog2(TIF) for azacytidine and Decitabine at all three concentrations with the 5mC z-score.

**Figure S5.**
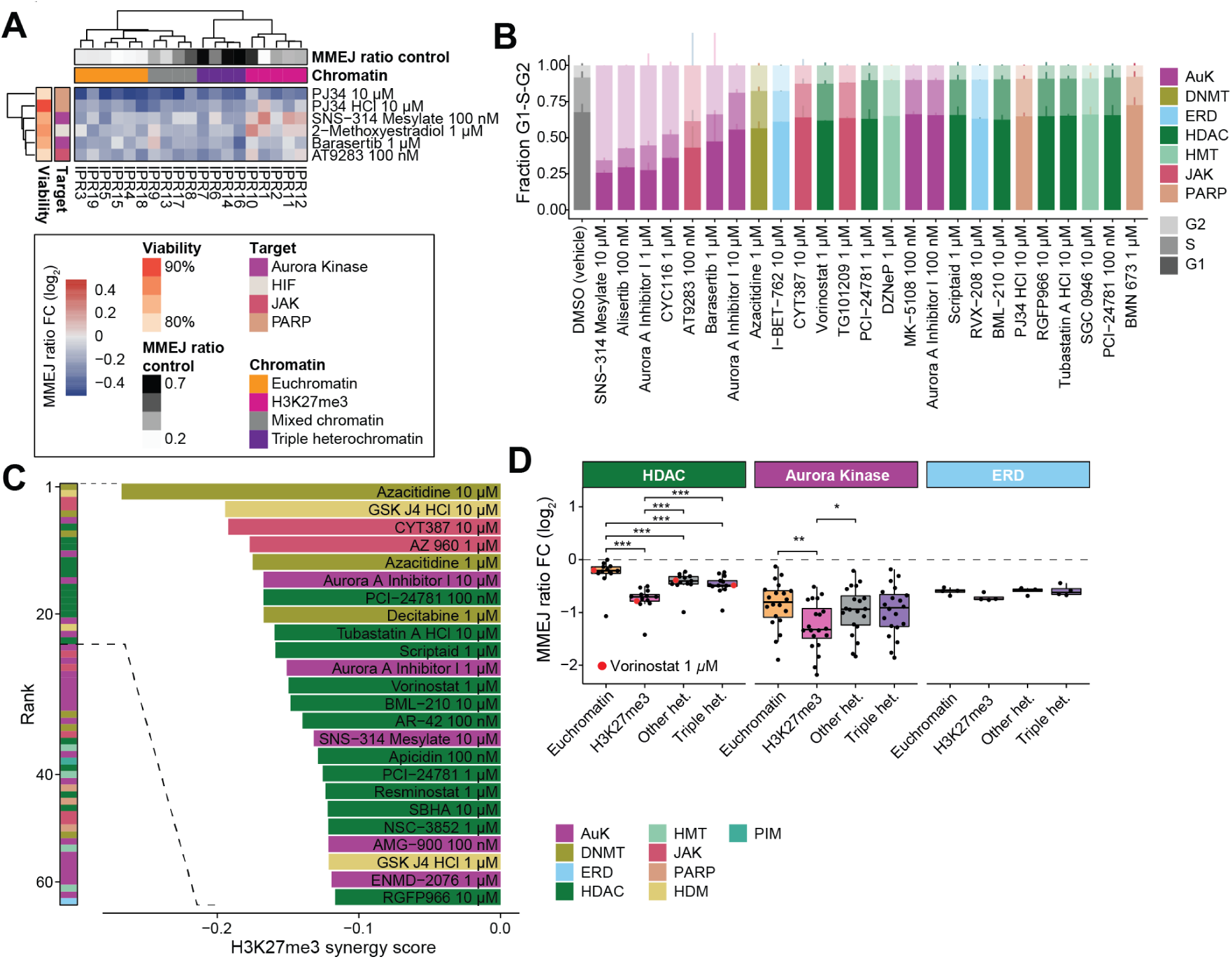
Details about global MMEJ:NHEJ ratio changes. (A) Heatmap showing the drugs with the most significant increases in MMEJ:NHEJ ratio per IPR. (B) Cell cycle plot displaying the fraction of cell cycle stages for 25 drug treatments and vehicle control (DMSO at equal volume). Mean and SD of two replicates is shown. (C) Bar plot showing the most significant synergies of the MMEJ:NHEJ ratio FC with H3K27me3 z-scores. The left panel shows the ranking colored by target and right panel shows the strongest 25 synergies. (D) Comparison of the MMEJ:NHEJ ratio FC per chromatin group between all significant drugs classified as ERD, aurora kinase or HDAC inhibitors. Each dot represents the MMEJ ratio FC for a drug in a given chromatin state. The red dots highlight the drug Vorinostat at 1 µM.

**Supplementary Table S1.** Detailed information about the drugs used in the screening. Abbreviations: HIF = hypoxia-inducible factor prolyl hydroxylase, DNMT = DNA-methyltransferase, HAT = histone acetyltransferase, PIM = proviral integration site for moloney murine leukemia virus kinase, HMT = histone methyltransferase, PARP = poly(ADP-ribose)-polymerase, JAK = Janus kinase, HDAC = histone deacetylase.

**Supplementary Table S2.**
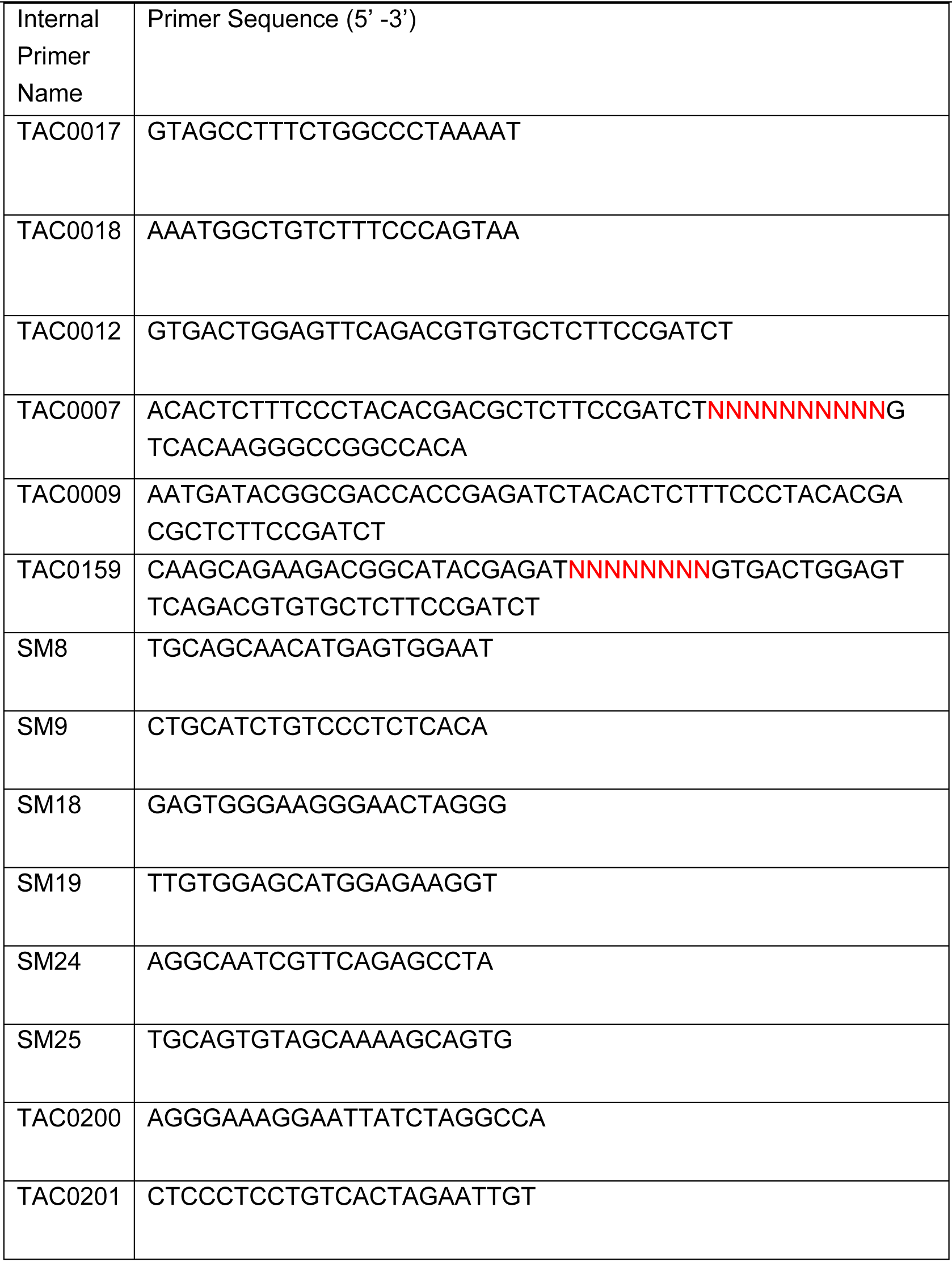
List of primers used.

**Supplementary Table S3.**
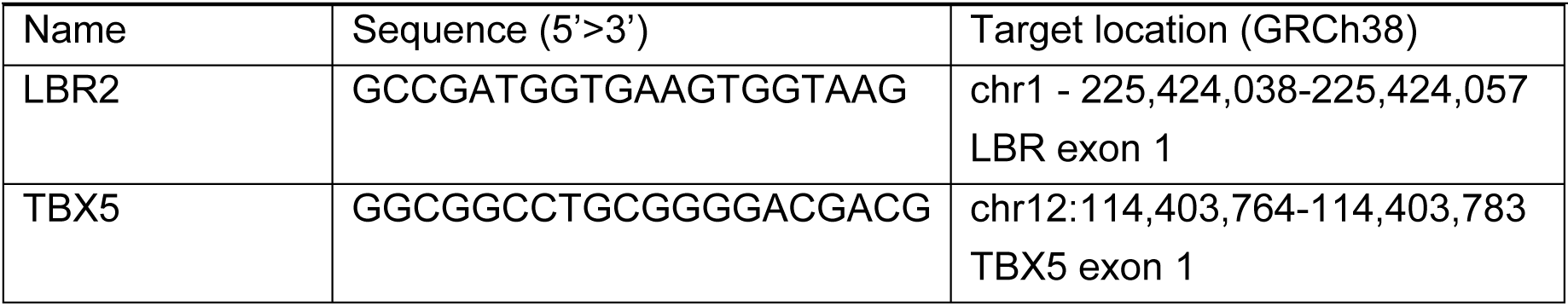
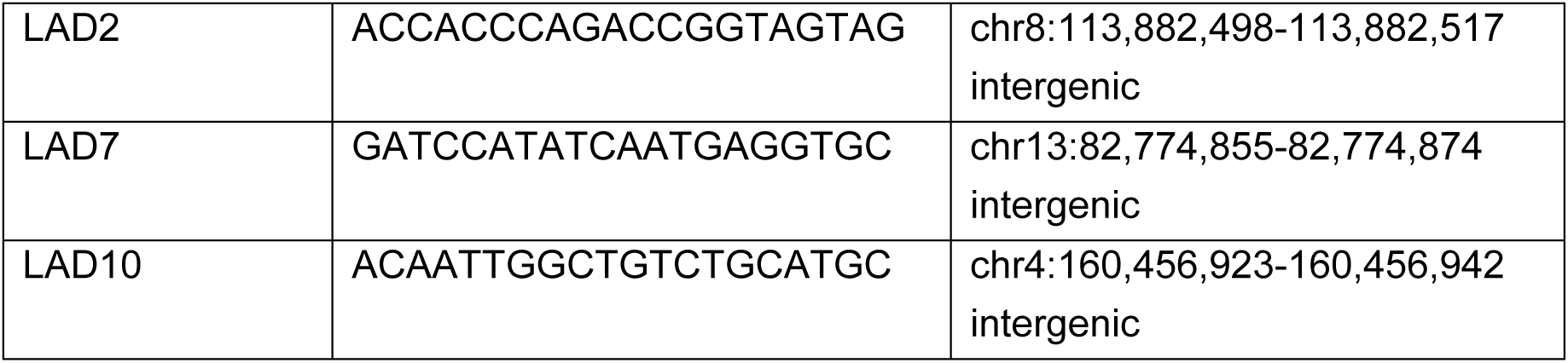
List of sgRNAs used.

